# *Polygonum Cillinerve* polysaccharide inhibits Transmissible gastroenteritis virus by regulating microRNA-181

**DOI:** 10.1101/2023.10.31.564899

**Authors:** Xueqin Duan, Huicong Li, Xuewen Tan, Nishang Liu, Xingchen Wang, Weimin Zhang, Yingqiu Liu, Wuren Ma, Yi Wu, Lin Ma, Yunpeng Fan

**Affiliations:** College of Veterinary Medicine, Northwest A&F University, 712100, Yangling, P R China; Institute of Traditional Chinese Veterinary Medicine, Northwest A&F University, 712100, Yangling, P R China; Agricultural Management Department, Sichuan Xuanhan Vocational Secondary School, 636350, Xuanhan, P R China; Nanjing Agricultural University, 210095, No 1 Weigang, Nanjing, China

**Keywords:** Transmissible gastroenteritis virus, miR-181, *Polygonum Cillinerve* polysaccharide, Apoptosis, Viral replication

## Abstract

Transmissible gastroenteritis virus (TGEV) is an important pathogen that can cause changes in the expression profile of cell miRNA. In this study, PCP could treat piglets infected with TGEV by in vivo experiment. High-throughput sequencing technology was used to detect that 9 miRNAs were up-regulated and 17 miRNAs were down-regulated during PCP inhibition of TGEV infection in PK15 cells. Meanwhile, miR-181 was related to the target genes of key proteins of apoptosis pathway. PK15 was treated with different PCP after transfection of miR-181 mimic or inhibitor. Real-time PCR was used to detect the effect of TGEV replication, electron microscopy, TEM and Hoechst fluorescence staining were used to detect the biological function of the treated cells, and western blot (WB) was used to detect the expression of key signaling factors cyt C, caspase 9 and P53 in the apoptotic signaling pathway. The results showed that compared with the control group, 250 μg/mL PCP significantly inhibited the replication of TGEV gRNA and gene N (*P* < 0.01). PK15 cells in PCP groups (250 and 125 μg/mL) had uniform cell morphology and a small number of floating cells by microscopic, no typical virus structure was observed in 250 μg/mL PCP group under TEM, and apoptosis staining showed that 250 μg/mL PCP significantly reduced the number of apoptotic cells. PCP may inhibit TGEV-induced apoptosis through Caspase-dependent mitochondrial pathway after transfection of miR-181. The above results provided a theoretical basis for further research on the mechanism of PCP anti-TGEV.

## 1. Introduction

Transmissible gastroenteritis (TGE) has been circulating in pigs for decades since its outbreak in the 1940s (Yan et al., 2022). The etiological agent is porcine transmissible gastroenteritis virus (TGEV). TGEV, a member of the coronavirus family, which belongs to the order Nestviridae, and can cause emeses, diarrhea, dehydration and high morbidity and mortality in pigs of all ages, especially newborn piglets (Zhang et al., 2014, Zhao et al., 2019). TGEV is enveloped, single-stranded, positive-sense RNA virus, and the viral genome consists of open reading frame (ORF) 1a (ORF1a), ORF1b, S, ORF3a, ORF3b, E, M, N, and ORF7a (Laude et al., 1990).

Currently, there are no specific drugs for the prevention and control of TGEV infection. At present, various anti-TGEV vaccines have been widely used in clinical practice. Unfortunately, the effect of vaccines is not rational. Therefore, drugs that can effectively prevent and/or treat TGEV infection need to be found. Antivirals are derived from organisms or synthesized chemically to tamper with one or more stages of the viral life cycle (including cell attachment, cell penetration, viral uncoating, viral genome (DNA/RNA) copy, maturation, and release of viral virion), and are important tools for improving vaccine action (Ali et al. 2021).

microRNAs (miRNAs) are the class of endogenous non-coding RNAs approximately 19-25 nt in length that have been identified as important cytoplasmic regulators of gene expression. It plays an important role in cell differentiation, development, cell cycle regulation and apoptosis (Felekkis et al. 2010). Most miRNAs have high sequence conserved expression sequence and tissue specificity. miRNA and its target genes have complex interactions, which depend on the following factors: (1) subcellular location of miRNA; (2) Abundance and target mRNA of miRNA; (3) miRNA-mRNA interaction affinity (Rani et al. 2022). A single miRNA can target hundreds of mRNAs, in contrast, an mRNA can be targeted by multiple miRNAs, and the many-to-many relationship between miRNA and mRNA leads to more complex miRNA regulatory mechanisms (Liu et al. 2014; Lu et al. 2018). Statistically, miRNAs control over 50% of mammalian protein-coding genes (Simonson et al. 2015). Cell free miRNAs have been reported to mediate short and long distance communication between various cells and may influence different physiological and pathological processes (Turchinovich et al. 2016). miRNAs have also been introduced as therapeutic agents or therapeutic targets to treat diseases.

Previous studies have demonstrated that *Polygonum Cillinerve* polysaccharide (PCP) possessed the better anti-TGEV effects in vitro (Pan et al. 2021), but the mechanism of action is still unknown. In order to further explore the antiviral mechanisms of PCP, in this study, the anti-TGEV effect of PCP in vivo was measured firstly; subsequently, miR-181 associated with the anti-TGEV effect of PCP was screened out and confirmed by high-throughput sequencing methods, then the effect of miR-181 in the process of PCP inhibiting TGEV was studied by real time PCR, fluorescence staining, transmission electron microscopy, and other methods. The aim was to further explore the mechanism of PCP against TGEV activity and provide the theoretical foundation for the development of antiviral.

## 2. Materials and methods

### 2.1. Cells culture and virus

PCP was prepared in our laboratory, which was prepared according to previous method (Zhou et al., 2019). Pig kidney (PK15) cells (collection number: GDC0061) were purchased from the Typical Culture Storage Center of Wuhan University. All cells were maintained in DMEM (Beijing Solebao Technology Co., Ltd., China) medium containing 10% FBS (SIGMA, Australia) at 37 ℃ in a 5% CO_2_ incubator. TGEV was donated by Dr. Wang from Henan University of Animal Husbandry and Economics. The infection dose of virus was measured was determined by MTT method (Ren et al., 2011).

### 2.2. Reagents

MTT was purchased from Beijing Solebao Technology Co., Ltd.; Lipofectamine 2000 transfection reagent was Thermo Fisher Scientific products; 2×Fast qPCR Master Mixture (Green) fluorescence quantitative kit, RNA extraction kit and reverse transcription kit were purchased from Beijing Dining Biotechnology Co., Ltd.; The primers were designed and synthesized by Shenggong Bioengineering (Shanghai) Co., Ltd. miR-181 mimic, miR-181 inhibitor, mimic negative control (mimic NC) and inhibitor negative control (inhibitor NC) synthesized by Guangzhou Ruibo Biotechnology Co., Ltd. Apoptosis-Hoechst staining kit was purchased from Beijing Suolaibao Technology Co., Ltd. P53, cyt C, caspase 9 and β-actin primary antibody, goat anti-rabbit IgG and goat anti-mouse IgG were CST Inc. products.

### 2.3. Animal and experimental design

All animal experimental procedures were performed in compliance with the Guideline for Ethical Review of Experimental Animal Welfare (GB/T 35892-2018), and the experiment was approved by the Animal Ethics and Welfare Committee of Northwest A&F University (DY2022022). A total of sixteen crossbred piglets (Duroc × Landrace × Large white) were weaned at 28-day old with an average body weight of 8.0 ± 1.4 kg. The piglets were kept in an air-conditioned nursery room with a temperature of 25–28 ℃ and a relative humidity of 50–60%.

After feeding for 5 days, 16 piglets were randomly divided into 2 groups, which were PCP group and virus control group. Then, all pigs were given 15 mL of TGEV orally for artificial modeling. After the piglets showed diarrhea and emesis, PCP was treated orally with 15 mL for each pig and 15 mL of normal saline for the virus group. The drug was administered continuously for 5 days. The symptoms of piglets were observed every day, and the dead pigs were dissected. Until the rest of the piglets returned to normal, the piglet survival rate was counted.

### 2.4. Effect of PCP on the expression profile of PK15 miRNAs infected with TGEV

#### 2.4.1. Sample preparation and sequencing

PK15 cells were seeded into cell dishes and cultured at 37 °C, 5% CO_2_ to about 80% cell abundance. Then the cells in experimental group were administrated with TGEV and 250 μg/mL PCP, and the virus control group was set. The cells were continued to cultivate for 24 h, and the cells were harvested and lysed with TRizol. Then Nanjing Personalbio Gene Technology Co., LTD. (Wuhan) was commissioned to conduct miRNA high-throughput sequencing to find differentially expressed miRNAs, and the experiment was repeated for three times.

#### 2.4.2. Target gene prediction and GO and KEGG analysis

This experiment used DESeq (version 1.18.0, Anders S and Huber W, 2010) to analyze the differences in miRNAs expression and filtered out the differentially conserved miRNAs based on |log2FoldChange|>1 and *P-value (P*)< 0.05. Target genes of miRNAs with significant differences were predicted by miRanda software. And then, Gene Ontology (GO) and Kyoto Encyclopedia of Genes and Genomes (KEGG) enrichment analysis were performed on differentially expressed genes to determine the biological functions or pathways that the differentially expressed genes are primarily affected.

#### 2.4.3. Verification of differentially expressed miRNAs

RNA was extracted according to the method of 2.4.1. Meanwhile, total RNA was reversely transcribed into cDNA by using 5×Integrated RT Master Mix Reverse Transcription Kit, reverse transcription primers of miR - 181 was GTCGTATCCAG TGCAGGGTCCGAGGTATTCGCACTGGATACGACAACTCA. Then, cDNA was amplified according to the 2×Fast Real time PCR Master Mixture (Green) kit. Primer of miR - 181 is F: CGAACATTCAACGCTGTCGG, R: AGTGCAGGGTCCGAGGT ATT. The reaction program was set according to the instructions. The results of Real-time PCR were analyzed by 2^-ΔΔCt^ method.

### 2.5. Verification of transfection effect

PK15 was placed in the 6-well cell plate, and when the cells grew to about 80%, miR-181 mimic or inhibitor and corresponding negative control (NC) were transfected into the cells according to the Lipofectamine 2000 kit instructions. After that, the expression level of miR-181 was verified according to the method of 2.4.3.

### 2.6. Detection TGEV genome and subgenome by Real-time PCR

PK15 were placed in the 6-well cell plate, and when the cells grew to 30%-40%, miR-181 mimic or inhibitor and corresponding negative control (NC) were transfected into the cells according to the Lipofectamine 2000 kit instructions. In the meantime, the corresponding mimic control or inhibitor control was set. After cultivated for 24 h, TGEV and different concentrations of PCP (250-62.5 μg/mL) were added. Then cells were cultivated for 24 h for subsequent experiments. According to the methods of 2.4.3, the effect of miRNA on TGEV genome and subgenome replication was analyzed by 2^-ΔΔCt^ method using β-actin as internal reference.

### 2.7. Sample Preparation and observation by TEM

After digestion with 0.25% trypsin containing EDTA, the cells were washed three times with PBS. Then cells were centrifuged at 1000 r/min for 5 min, and supernatant was discarded. Cells were fixed in 2.5% glutaraldehyde for 4 h, followed by 1% osmic acid for 2 h. It was dehydrated with gradient ethanol, then embedded with epoxy resin and placed in a drying oven. Cell lumps were trimmed and sectioned and stained with uranium. Finally, the transmission electron microscope (TEM) was used to observe and photograph.

### 2.8. Measurement of apoptosis by Hoechst 33258 staining

The cells were stained according to the instructions of Apoptosis-Hoechst staining kit. Finally, cells were observed and photographed under the fluorescence microscope.

### 2.9. Measuring the protein expression of cyt C, caspase 9, P53 by western blot

According to the methods of 2.6, the protein were extracted by the protein extraction kit. Then BCA quantification kit was used for protein quantification, and 10% and 12% separation gels were prepared. 20 μL of protein was added to each well of the gel. Denatured proteins were separated by SDS-PAGE (80V, 20 min; 120 V, 80 min) and transferred to PVDF membrane (100 V, 66 min). The membrane was blocked by TBST contained 5% skimmed milk at 4 ℃ for the night, and then incubated overnight in TBST contained specific primary antibodies at 4℃. After washed 4 times with TBST, the membrane was incubated with the specific secondary antibody for 1 h. Finally, the membrane was washed 4 times with TBST, the signal was observed with ECL, and the membrane was exposed on the X-ray film.

### 2.10. Statistical analysis

Statistical software IBM SPSS Statistics 21.0 was used to analyze the data. The data were expressed as Mean ± SD. Comparison between groups was performed using Oneway-anova statistical method. *P*<0.05 was statistically significant *P*\*<0.05, *P*\*\*<0.01, *P*\*\*\*<0.001.

## 3. Results

### 3.1. Therapeutic effect was established by PCP on TGEV-infected piglets

After 5 days of modeling, the piglets were depressed and showed symptoms such as diarrhea and emesis (Fig.1A and B). Autopsy of dead piglets showed that the intestines of infected pigs were full of gas, the tube walls were thinner and transparent or translucent (Fig.1C), and the intestinal contents were yellow and foamy (Fig.1D). In PCP group, all piglets returned to normal and their body weight increased gradually (Fig. 2A) on day 9 after the first administration. A total of 4 piglets died in this group (Fig. 2B), with the mortality rate of 50% (Fig. 2C) and the cure rate of 50% (Fig. 2D). All piglets in the TGEV group died on day 10 (Fig. 2B), with the mortality rate of 100% (Fig. 2C). The results showed that PCP had a good therapeutic effect on piglets infected with TGEV.

**Fig. 1.**
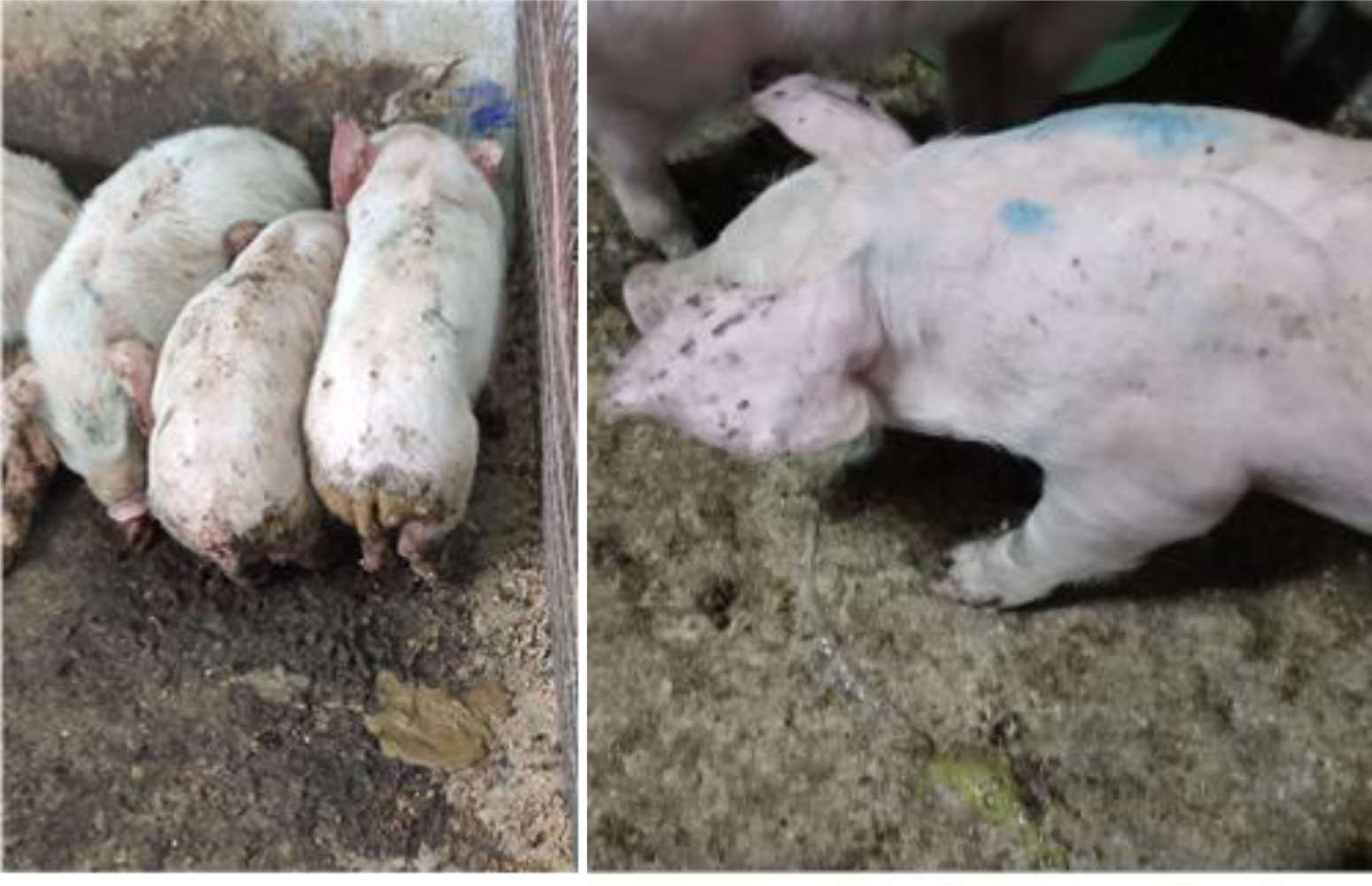
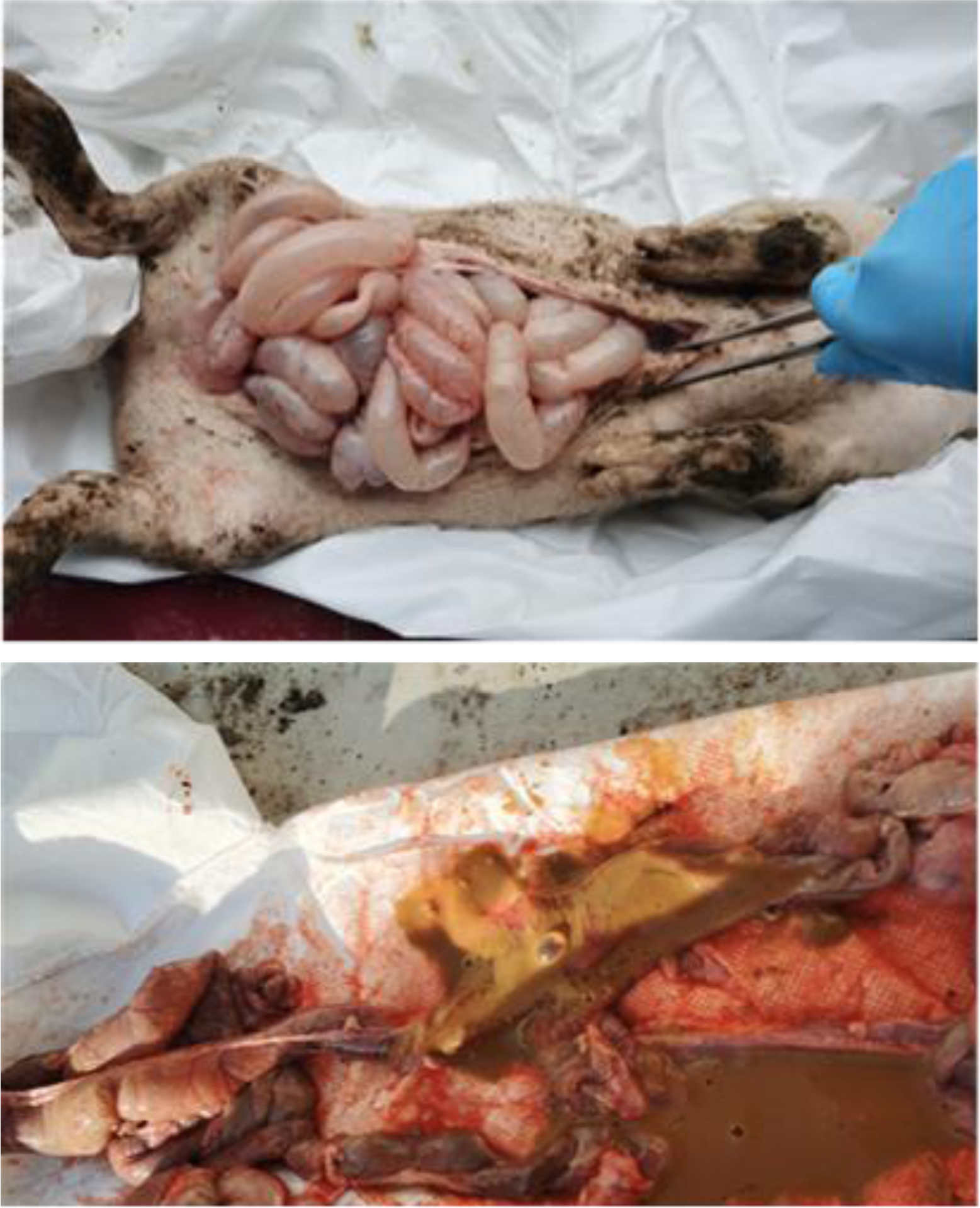
Clinical symptoms and pathological features of piglets. A: Piglets infected with TGEV developed diarrhea, B: piglets infected with TGEV developed vomiting, C: The intestines of the dead piglets were full of gas, D: the intestines of the dead piglets had yellow contents.

**Fig. 2.** Body weight, mortality and cure rate of piglets. A: Change in piglet weight, B: Number of piglet deaths, C: Piglet mortality, D: Piglet cure rate.

### 3.2. High-throughput sequencing analysis

To investigate the mechanism of PCP inhibiting TGEV to infect PK15 cells, the high-throughput sequencing was used to investigate the changes of miRNA expression profiles. Finally, 9 expressed differentially miRNAs were up-regulated and 17 expressed differentially miRNAs were down-regulated. The target genes of the differentially expressed miRNAs were predicted, and 12582 target genes were obtained. Cluster analysis was used to determine the expression patterns of differentially expressed miRNAs under different experimental conditions (Fig. 3). By GO pathway analysis, the result showed that binding, metabotic process and nitrogen compound metabotic process were significantly enriched after virus infection (Fig. 4A). By KEGG pathway analysis, the result showed that several signal pathways were significantly enriched after virus infection, such as Endocytosis, MAPK signaling pathway and RAS signaling pathway (Fig. 4B). Considering the important role of TGEV-induced apoptosis in host response to TGEV infection (Wang et al. 2022), the expression patterns of differentially expressed miRNAs involved in apoptosis were analyzed. Among them, the expression of miR-181 markedly reduced after PCP and TGEV infection, and the result was also confirmed by Real-time PCR (*P <* 0.001) (Fig. 5A), suggesting that miR-181 might be involved in PCP anti-TGEV-induced apoptosis. As it was reported that miR-181 was involved in the regulation of tumor cell apoptosis (Huang et al. 2015), therefore, miR-181was screened out to examine the role in PCP response to TGEV infection.

**Fig. 3.**
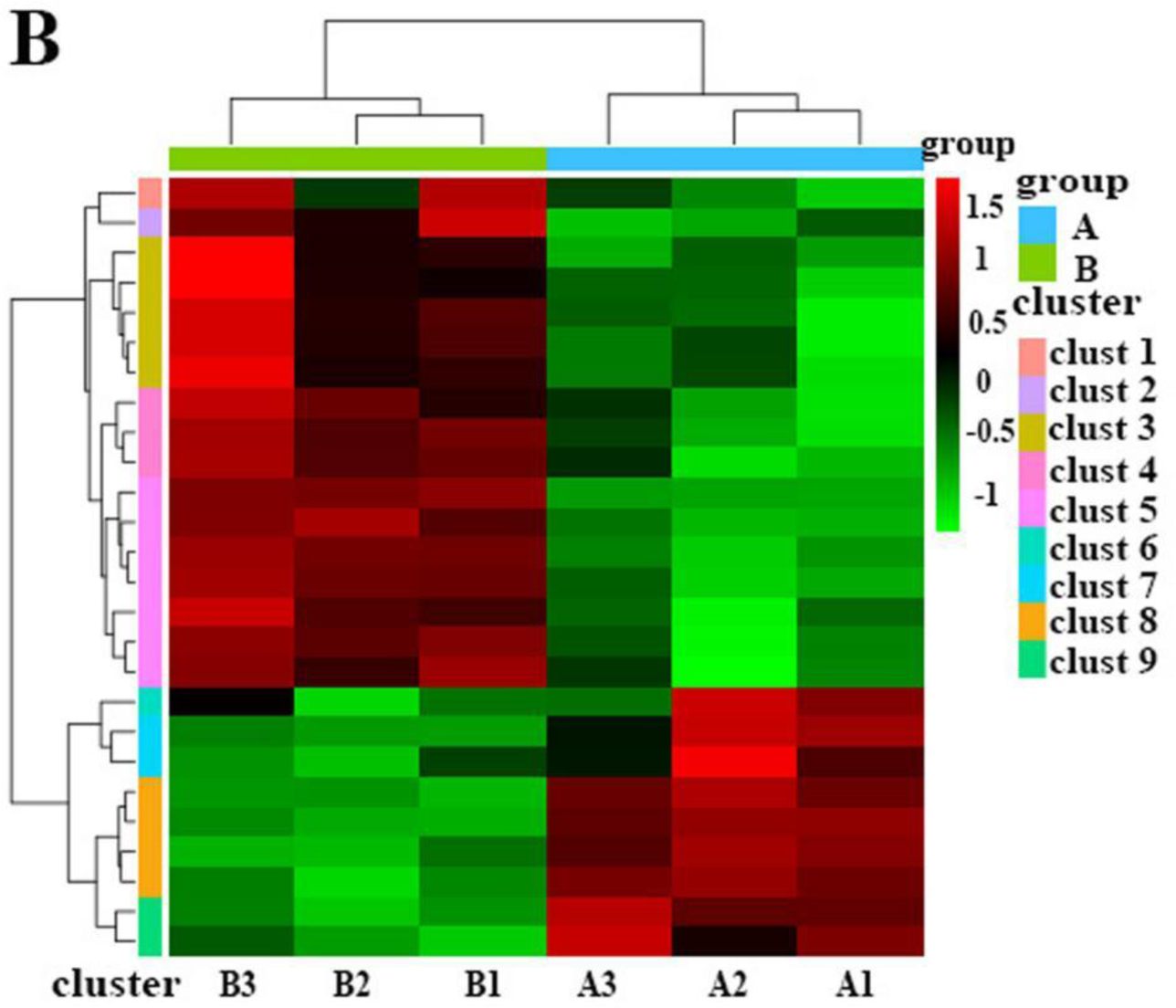
High-throughput sequencing analysis. A: 9 up-regulated miRNAs and 17 down-regulated miRNAs, B: The expression patterns of differentially expressed mirnas under different experimental conditions were determined by cluster analysis.

**Fig. 4.**
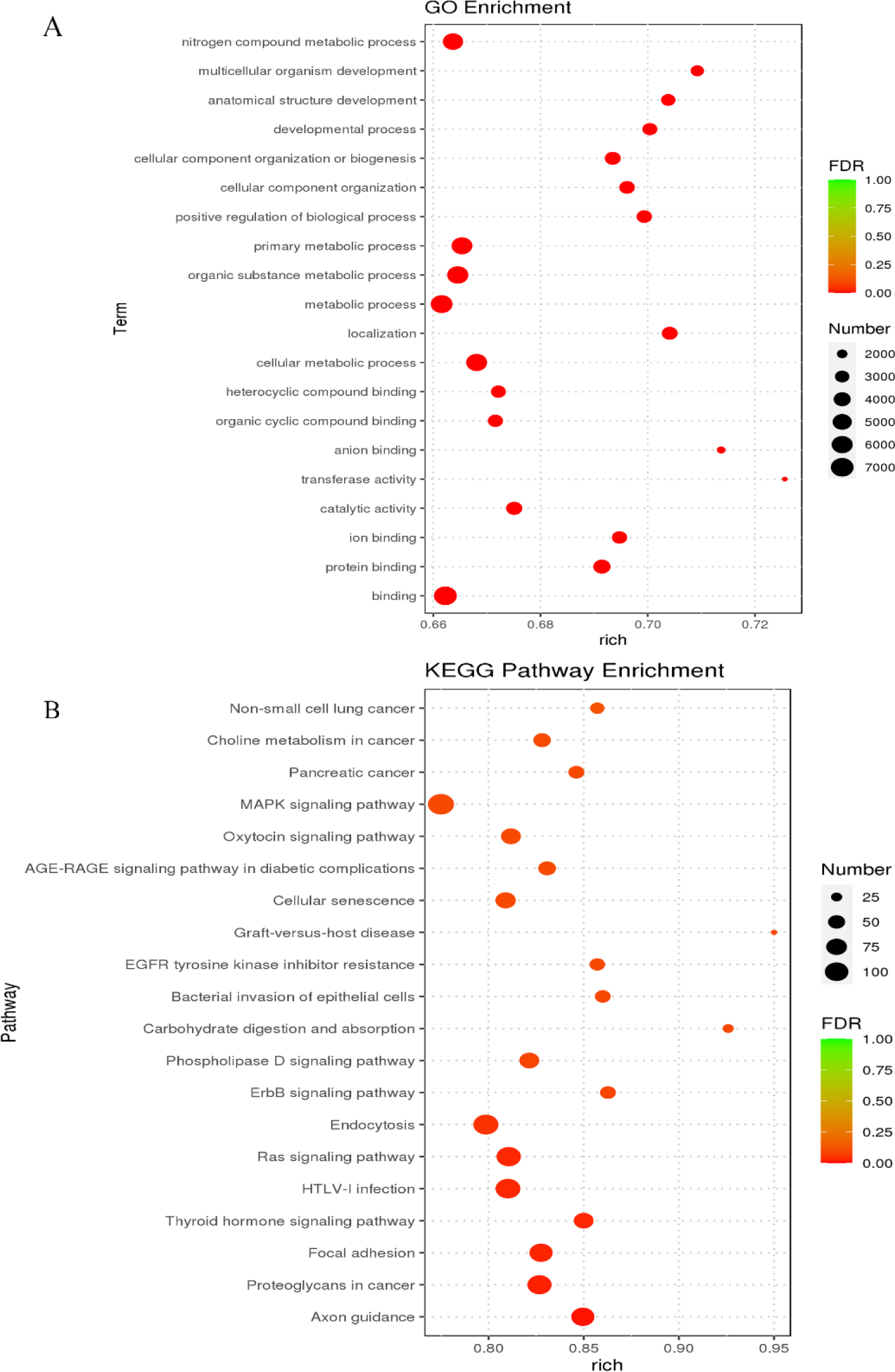
GO and KEGG enrichment analysis bubble map. A: GO enrichment analysis bubble map B: KEGG enrichment analysis bubble map.

**Fig. 5.**
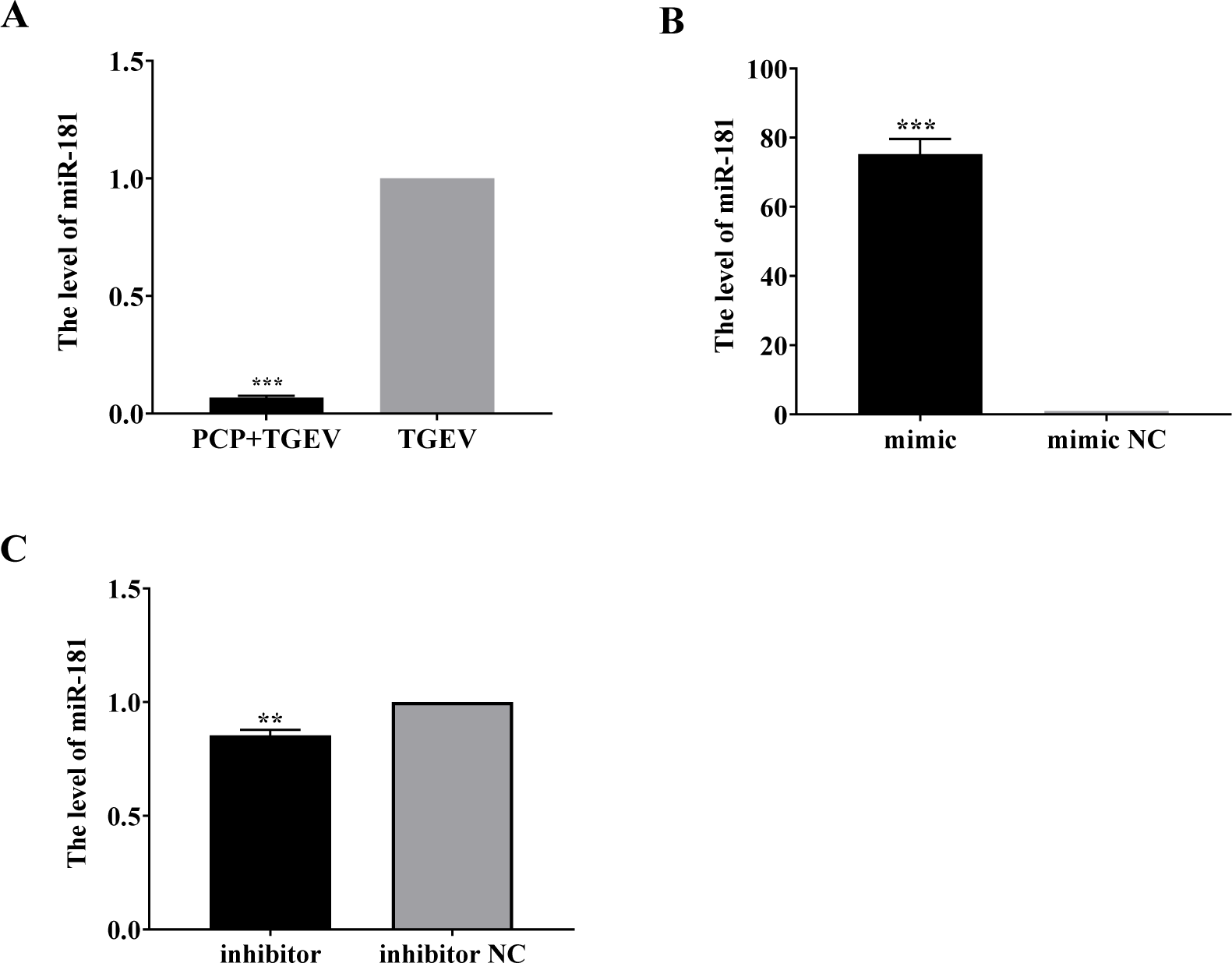
The level of miR-181. A: The level of miR-181 was verified by Real-time PCR, B: The level of miR-181 after transfection with mimic, C: The level of miR-181 after transfection with inhibitor

### 3.3. Transfection verification

The mimic and inhibitor of miR-181 were synthesized, and the changes were measured by real time PCR. The results showed that the expression of miR-181 was significantly up-regulated in PK15 cells after transfection with mimic (*P* < 0.001) (Fig. 5B). And the expression of miR-181 was considerably down-regulated following transfection with the inhibitor (*P* < 0.01) (Fig. 5C).

### 3.4. Effects of miR-181 on genomic and subgenomic of TGEV

After transfecting miR-181 mimic, the relative levels of TGEV gRNA in the mimic NC group were significantly lower than those in the mimic control group (*P* < 0.01) (Fig. 6A); The relative levels of TGEV gene N were significantly lower than those of mimic control group (*P* < 0.001) (Fig. 6B); Compared with mimic control group, PCP significantly inhibited the replication of TGEV gRNA and gene N at concentrations of 250-62.5 μg/mL (*P* < 0.001) (Fig. 6A and B). After transfection with miR-181 inhibitor, the relative level of TGEV gene N in inhibitor NC group was significantly higher than that in inhibitor control group (*P* < 0.001) (Fig. 6D); There were no significant differences in TGEV gRNA levels among the groups (Fig. 6C). Compared with inhibitor control group, PCP significantly inhibited TGEV gRNA replication at 250-62.5 μg/mL (*P* < 0.01) (Fig. 6C), PCP significantly inhibited the replication of TGEV gene N at the concentrations of 250 and 125 μg/mL (*P* < 0.001), PCP at 62.5 μg/mL significantly inhibited the replication of TGEV gene N (*P* < 0.01) (Fig. 6D). The results showed that miR-181 mimic promoted the replication of TGEV gRNA and gene N, while miR-181 inhibitor inhibited the replication of TGEV gene N. Meanwhile, PCP inhibited the replication of TGEV gRNA and gene N after transfection of miR-181 mimic and miR-181 inhibitor.

**Fig. 6.**
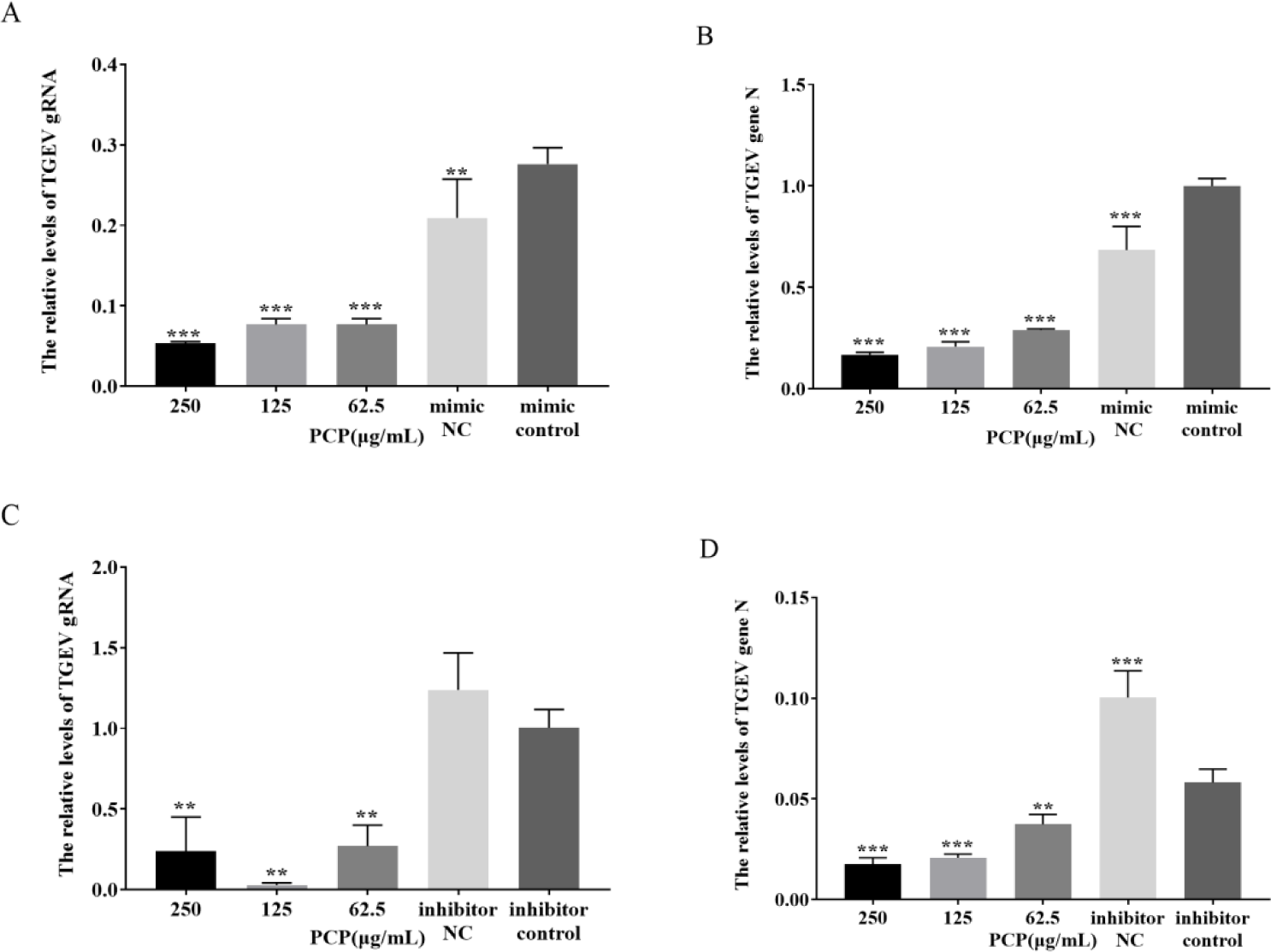
The level of TGEV gRNA and gene N. A: The level of TGEV gRNA after transfection with miR-181 mimic, B: The level of TGEV gRNA after transfection with miR-181 inhibitor, C: The level of TGEV gene N after transfection with miR-181 mimic, D: The level of TGEV gene N after transfection with miR-181 inhibitor.

### 3.5. Effect of transfection with miR-181 mimic or inhibitor on apoptosis

When apoptosis occurred, the cell apoptosis could be observed under fluorescence microscope after Hoechst 33258 staining. After transfecting miR-181 mimic, the nuclei of miR-181 mimic control group were more white than those of mimic NC group, so more apoptotic cells were found; The cells in 250 μg/mL group, 125μg/mL group and Blank group were in good condition with no obvious nuclear whitening, indicating no apoptosis (Fig. 7A). After transfection with miR-181 inhibitor, there was no difference between inhibitor NC group and inhibitor control group. In Blank group, the cell state was the best and there was no obvious nuclear whitening, basically no cell apoptosis. There were fewer apoptotic cells in 250-62.5 μg/mL group compared with inhibitor control group (Fig. 7B).

**Fig. 7.**
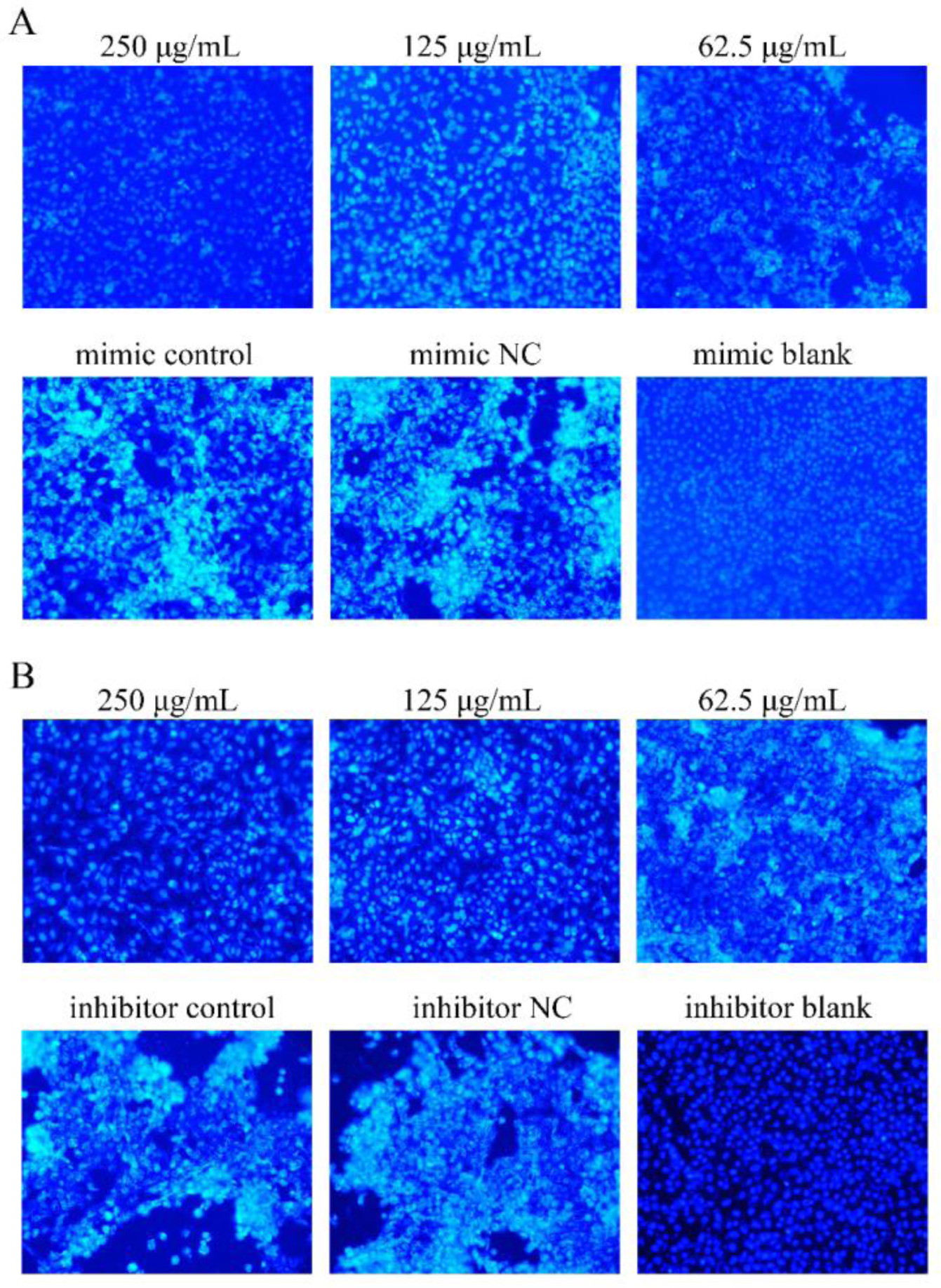
Detection of cell apoptosis by Hoechst 33258 (200×) A: Apoptosis after transfection with miR-181mimic, B: Apoptosis after transfection with miR-181inhibitor

### 3.6. Observed cell changes by transmission electron microscopy

After transfection with mimic, no typical virus structure was showed in Fig. 8A and B, and virions are showed in Fig. 8C-F. In Fig. 8C and D, virions are mostly clustered and distributed in the cytoplasm, wrapped by vesicles, and a small number of virions can be seen around the nucleus and nuclear membrane. In Fig. 8E and F, virions were mostly distributed in the nucleus, clustered around the nuclear membrane, and the small amount scattered in the cytoplasm.

**Fig. 8.**
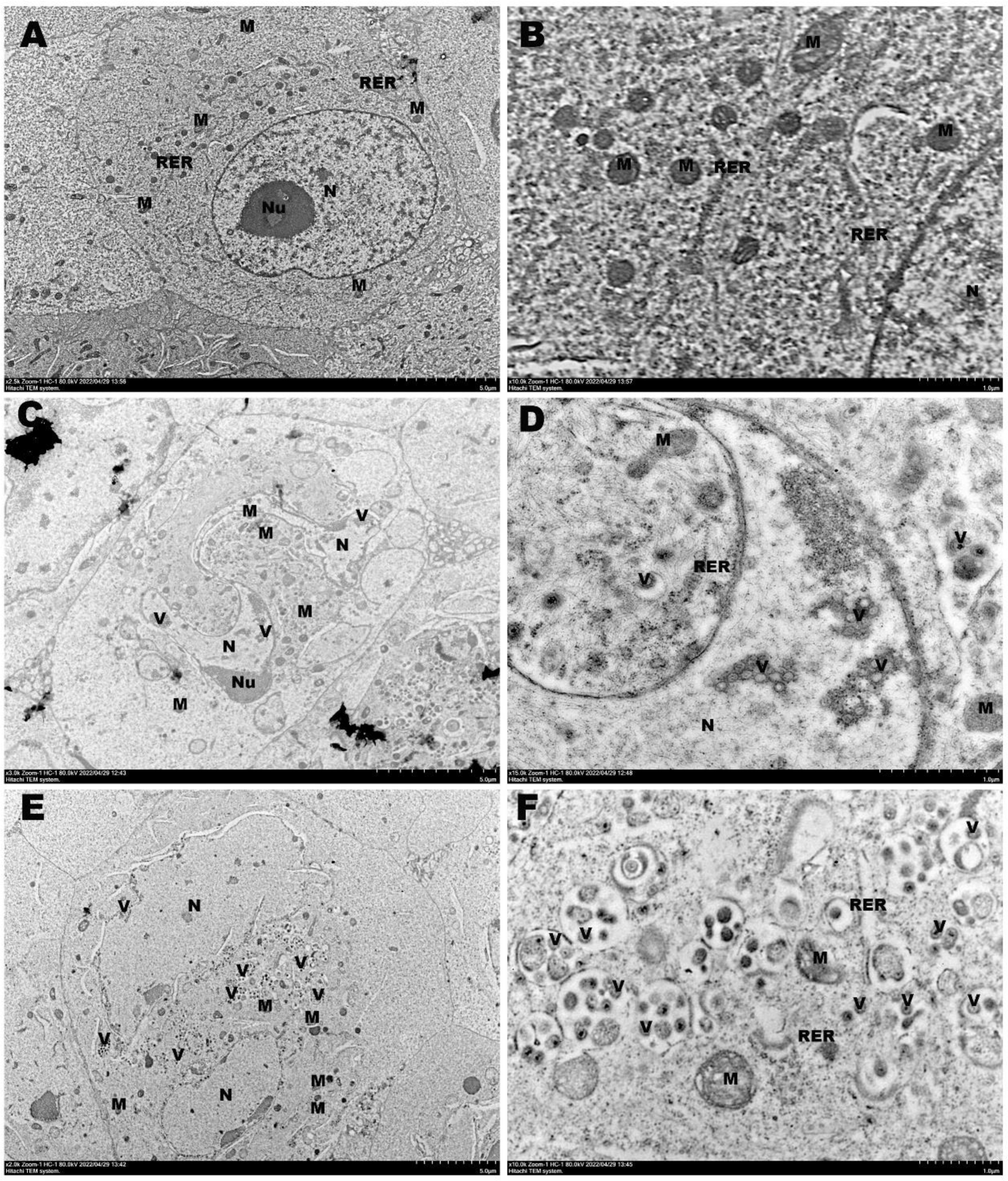
Electron micrograph of TGEV and cell after transfection with miR-181 mimic. A: Electron micrograph of the drug group (×2.5 k); B: Electron micrograph of the drug group (×10.0 k); C: Electron micrograph of NC group (×3.0 k); D: Electron micrograph of NC group (×15.0 k); E: Electron micrograph of control group (×2.0 k); F: Electron micrograph of control group (×10.0 k). Note: mitochondria (M); nucleus (N); nucleoli (Nu); rough endoplasmic reticulum (RER); viral particles (V).

After transfection with inhibitor, there were some differences among the three groups of cells. Mitochondria in Fig. 9C and D had the most serious damage, showing severe swelling, massive dissolution of part of the matrix and vacuolation. Fig. 9A and B showed mild to moderate swelling with shallow and dissolved stroma. No obvious swelling was observed in mitochondria of Fig. 9E and F, the electron density of matrix increased significantly, and a few cristae expanded slightly. Rough endoplasmic reticulum of Fig. 9A and B with mild dilatation and local degranulation; No significant swelling was seen in the other two groups. A large number of virions could be seen in the cells and organelles of Fig. 9C and D; more virions could be seen in the nuclei of Fig. 9E and F; no typical virions were seen in Fig. 9A and B.

**Fig. 9.**
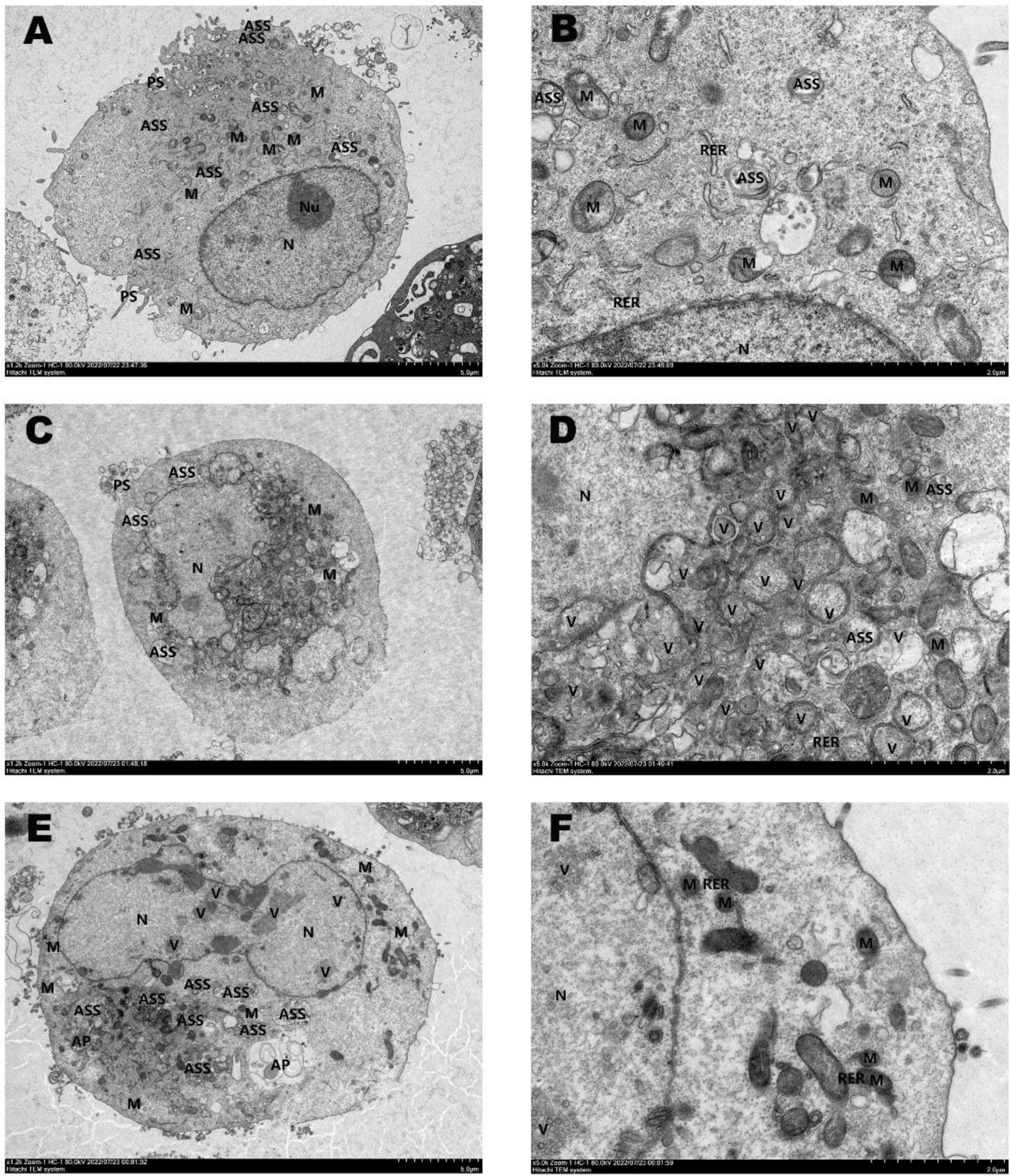
Electron micrograph of TGEV and cell after transfection with miR-181 inhibitor. A: Electron micrograph of the drug group (×1.2 k); B: Electron micrograph of the drug group (×5.0 k); C: Electron micrograph of NC group (×1.2 k); D: Electron micrograph of NC group (×5.0 k); E: Electron micrograph of control group (×1.2 k); F: Electron micrograph of control group (×5.0 k). Note: feet (PS); mitochondria (M); nucleus (N); nucleoli (Nu); rough endoplasmic reticulum (RER); autolysosome (ASS); viral particles (V).

### 3.7. Effects of transfection of miR-181 mimic or miR-181 inhibitor on the expression of key signaling molecules

The mimic inhibited the protein expression of cyt C (Fig. 10B) (*P* < 0.05) after transfecting with miR-181 mimic. Compared with mimic control, the protein expression of cyt C was significantly inhibited when PCP concentration was 250-62.5 μg/mL (Fig. 10B) (*P* < 0.001); PCP concentration of 250 μg/mL significantly increased the protein expression of cleaved caspase 9 (*P* < 0.01) (Figure 10C). After transfection with miR-181 inhibitor, protein expression of cyt C, cleaved caspase 9, and P53 (Fig. 11B-D) in inhibitor control group was not significantly different from that of mimic NC group. Compared with inhibitor control, PCP concentration of 250 μg/mL significantly inhibited the expression of cyt C and P53 proteins (Fig. 11B and D) (*P <* 0.05). Therefore, miR-181 mimic inhibited the protein expression of cyt C. Meanwhile, PCP inhibited the expression of cyt C and caspase 9 in PK15 cells infected with TGEV after transfection of miR-181 mimic, and inhibited the expression of cyt C and P53 in PK15 cells infected with TGEV after transfection of miR-181 inhibitor.

**Fig. 10.**
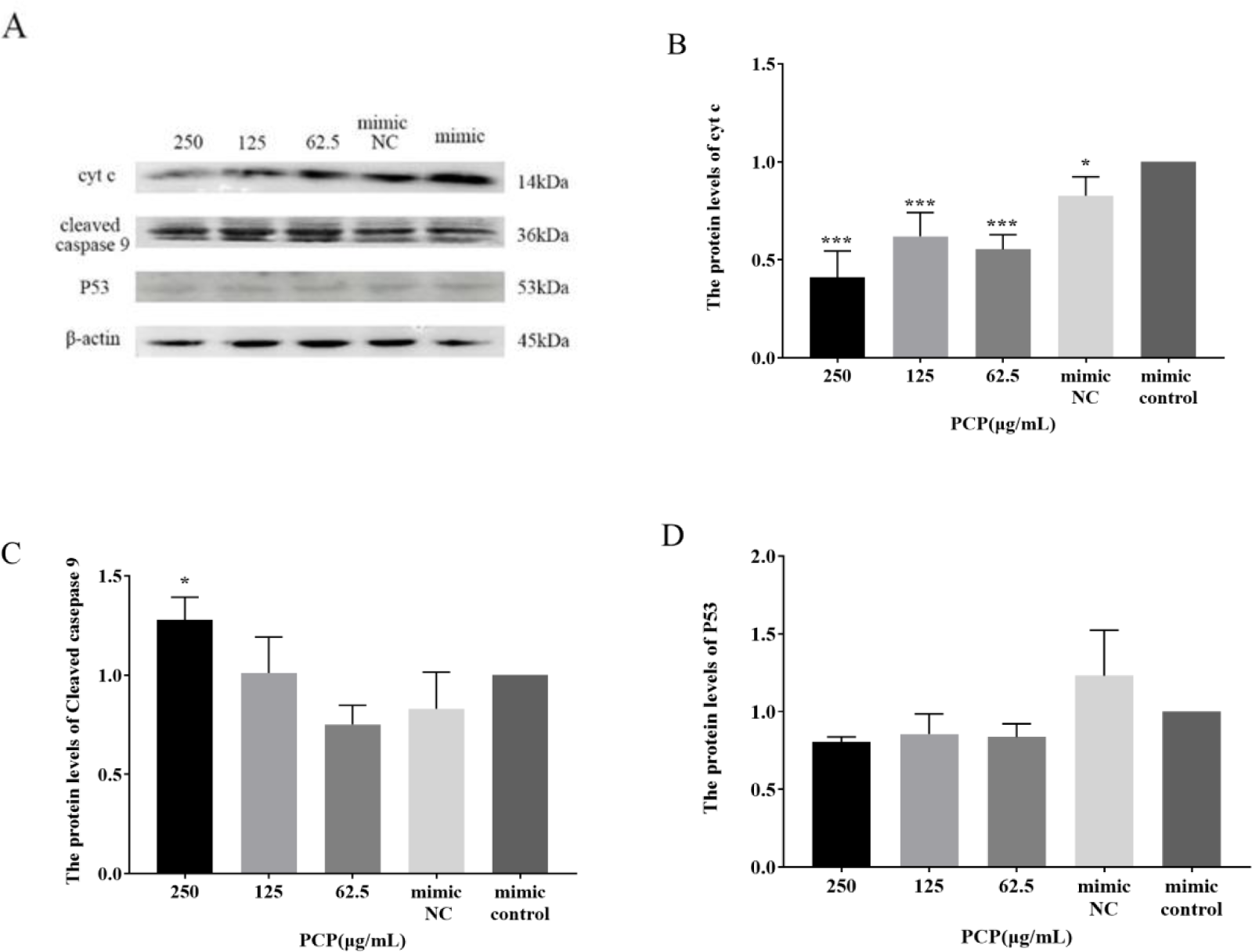
Effect of transfected miR-181 mimic on protein expression in cells. A: Protein bands; B: The protein expression levels of cyt C; C: The protein expression levels of cleaved caspase 9; D: The protein expression levels of P53

**Fig. 11.**
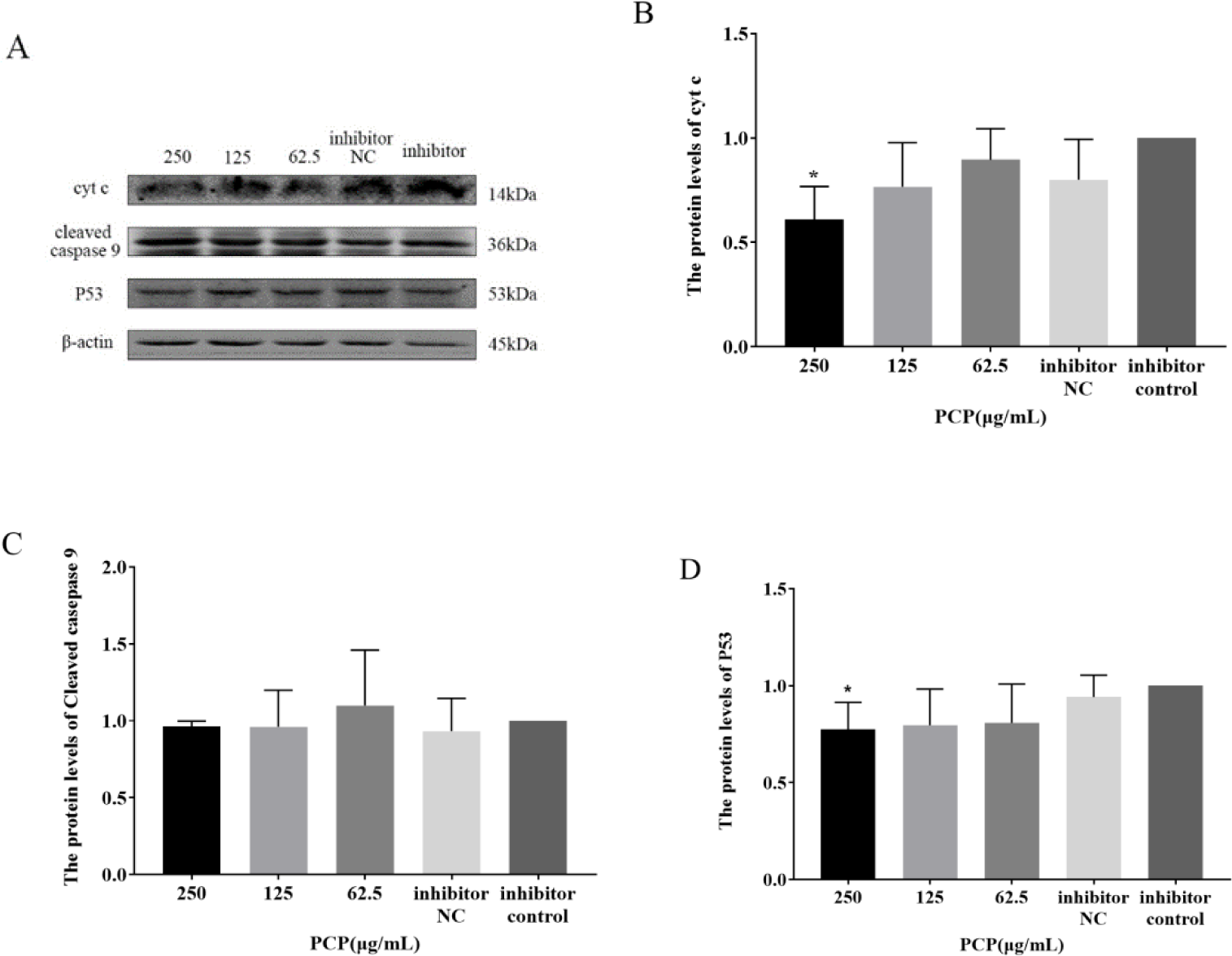
Effect of transfected miR-181 inhibitor on protein expression in cells. A: Protein bands; B: The protein expression levels of cyt C; C: The protein expression levels of cleaved caspase 9; D: The protein expression levels of P53

## 4. Discusion

At present, TGEV, as one of the porcine intestinal coronaviruses (PECs), is causing huge economic losses to the pig industry in China and the world, and urgently needs to be addressed. Due to the low efficacy, cytotoxicity, and viral resistance of synthetic antiviral drugs such as Moroxidine, Ribavirin, and Ganciclovir (Märtson et al. 2022, Yu et al. 2016), natural antiviral drugs are considered easy to accept because of their low toxicity and low price (Ghosh et al. 2009). PCP has been proven to have pharmacological effects such as antiviral, antibacterial, anti-inflammatory, and anti-tumor effects (Wang et al. 2022). Therefore, we conducted TGEV challenge on weaned piglets and then treated them with PCP. The results showed that PCP had a good therapeutic effect on piglets infected with TGEV. In our experiment, it was found that the jejunal wall of the piglets became thinner and the intestinal cavity expanded through dissecting piglets infected with TGEV and dying, which is consistent with the research results of Luo (Luo et al. 2019).

Since DNA sequencing technology opened the door to the era of gene research, the sequencing of the first human genome map was completed in 2001. However, due to the drawbacks of low throughput, high cost, and slow speed of the first generation sequencing technology, the next generation sequencing technology emerged as a revolutionary improvement on traditional sequencing technology (Lv et al. 2021). Therefore, as one of the important members of biological detection technology, high-throughput sequencing technology has the advantages of fast detection speed, high precision, low cost, wide coverage and large output (Schuster et al. 2008). Because the virus completely depends on the host cell for reproduction, it will change the microenvironment in the host cell (Duan et al. 2019); Meanwhile, miRNA is a biomarker of viral infection. Therefore, the miRNA expression profile undergoes changes due to viral infection, resulting in differential miRNA expression that can be attributed to host antiviral defense and changes in the cellular environment influenced by viral factors (Powdrill et al. 2016, Nejad et al. 2018, Dong et al. 2022).

Previously, 59 unique miRNAs showed significant differential expression between normal and TGEV infected ST cell samples, with 15 miRNAs significantly upregulated and 44 significantly downregulated (Liu et al. 2015); After TGEV infection with PK15 cells, there were 21 differentially expressed miRNAs, of which 13 were upregulated and 8 were downregulated (Song et al, 2016). The first reason for this difference can be considered as the different cells infected with TGEV, where the same stimulation can lead to different changes in the intracellular environment within different cells. Just like the results of GO enriched CC entries, the target genes for differentially expressed miRNAs were significantly enriched in the intraellarpart. We conducted high-throughput sequencing according to the screening results, and the results showed that 26 differentially expressed miRNAs were obtained after PCP was used, of which 9 miRNAs were up-regulated and 17 were down regulated. In the case of infecting the same cells, the results showed several identical differentially expressed miRNAs compared to Song, indicating that TGEV infection may affect changes in the expression of some identical miRNAs (Song et al, 2016). However, due to different detection techniques and treatment methods, the trend of these miRNAs is not completely consistent. This indicates that PCP has an impact on the miRNA expression profile of PK15 cells infected with TGEV.

The entry of coronavirus into cells depends on the binding of virus spike (S) protein to cell receptors and the activation of host cell protease S protein (Hoffmann et al. 2020). In GO enrichment analysis, the target genes are mainly enriched in binding and metabolism related processes, which is likely related to the virus entering cells. After TGEV enters cells, it mainly affects cell metabolism and other processes, facilitating the virus to control cell metabolism to replicate and evade host immune responses (Moreno-Altamirano et al. 2019). Due to the induction of cell apoptosis by coronavirus infection (Li et al. 2021), the expression pattern of differentially expressed miRNAs involved in PCP against TGEV induced PK15 cell apoptosis was analyzed and the validation was performed by real-time PCR. The results showed that the trend of changes in miR-181 was consistent with the sequencing results.

miRNAs in cells are involved in the life cycle of many viruses, and their importance in virus host interactions has also become increasingly apparent apart from their crucial role in regulating normal cell gene expression (Otsuka et al. 2007, Ahluwalia et al. 2008, Lecellier et al. 2005). For example, it was found that inhibition of miR-199 reduces HCV RNA replication, and Castillo’s study demonstrated that overexpression of miR-133 impairs DENV-2 replication, affecting the percentage of infected cells and the number of viral RNA copies produced (Wang et al. 2015, Castillo et al. 2016). In this study, the overexpression of miR-181 could inhibit the replication of TGEV, which was similar to the result of Castillo et al. Previously, someone studied that miR-4331 inhibits the transcription of TGEV gene 7 in PK15 cells through the target gene CDCA7 (Song et al. 2016), demonstrating that host miRNA can affect TGEV replication. The experimental results in this study also provided a reference for this viewpoint, as miR-181 affects the replication of gene N and TGEV gRNA.

Plants, as a source of natural compounds with medicinal importance, play an important role in the medical world, often providing important sources for the development of new antiviral drugs. The characteristics of these naturally sourced antiviral drugs reveal their interrelationships with the virus replication cycle, such as virus entry, replication, assembly, and release, as well as targeted precise virus host interactions (Ali et al. 2021). Here, we investigated the effect of PCP on TGEV replication, and our results showed that PCP inhibited the replication of TGEV gRNA and geneN after transfection with miR-181, similar to the significantly reduced TGEVN gene expression in PK15 cells treated with Astragalus polysaccharides and Bacillus subtilis metabolites (Wang et al., 2017). In summary, this study suggests that miR-181 affects the replication of TGEV in PK15 cells, and PCP may inhibit the replication of TGEV after transfection by downregulating the expression of miR-181. These findings further emphasize the important role of mammalian miRNAs in antiviral responses and enrich the antiviral mechanisms of traditional Chinese medicine, which may have important implications for the molecular mechanisms of miRNAs in the anti TGEV process of traditional Chinese medicine.

Transmission electron microscopy (TEM) is an ideal device for studying the internal structure of cells and different types of biomaterials. It is a valuable technique for imaging the ultrastructure of samples and determining virus host interactions at the cellular level (Doyle et al. 2020). Through the transmission electron microscopy, Song et al. observed that TGEV virus particles are roughly spherical in shape, with a diameter between 80 and 120 nm (Song et al. 2016), which is similar to our electron microscopy results, with suspected virus particles having a diameter between 80-140nm. We also observed that a large number of virus particles were gathered in the cells and organelle of the transfected inhibitorNC group, and more virus particles were seen in the nuclei of the transfected inhibitorNC group.

Hoechst33258 is a blue fluorescent dye that can penetrate cell membranes and emit strong blue fluorescence after embedding double stranded DNA. It has low toxicity to cells and is widely used to evaluate cell cycle and apoptosis (Zhang et al. 1999). In the process of apoptosis, chromatin coagulated white under fluorescence microscope. Research has shown that traditional Chinese medicine formulas or effective ingredients can regulate cell apoptosis by regulating miRNA. For example, Ditan Huoxue Shubi Tang (DHS) could upregulate the expression level of miR-148, downregulate the expression level of TXNIP, thereby reducing the autophagy and apoptosis levels of myocardial cells caused by H/R (Wang Shen 2022). Platycodon grandiflorum polysaccharides could improve epithelial cell apoptosis and inflammation induced by respiratory syncytial virus through activating miR-181 mediated Hippo and SIRT1 pathways (Li et al. 2022). This is similar to our results. After transfection with miR-181, the PCP group cells exhibited nuclear bleaching in a dose-dependent manner with drug concentration, indicating that high concentrations of drugs inhibited cell apoptosis. Therefore, it indicated that PCP could inhibit TGEV induced cell apoptosis by downregulating the expression of miR-181.

The miR-181 family is involved in a variety of cellular functions in normal cells, such as proliferation, growth, survival, cell death, and tumor suppression. In members of the miRNA-181 family, miR-181 could penetrate the mitochondria, translocation into the mitochondria to function, and the rest can function in the cytoplasm (Duarte et al. 2014). Although miR-181 has been shown to inhibit apoptosis through overexpression of Bcl-2 in cells (Zhu et al. 2013), in our prediction, it could regulate apoptosis in PCP - and TGEV-infected cells through cyt C. At the protein level, miR-181 mimic significantly inhibited the protein expression of cyt C (Fig. 10). However, interestingly, we found that miR-181 inhibitor showed a tendency to inhibit protein expression of P53, while miR-181 mimic showed a tendency to promote P53 protein expression. This is similar to how downregulation of miR-181 is associated with the accumulation of P53 in cancer (Rezaei et al. 2020). Our results also showed that PCP could significantly inhibit the protein expression of cyt C and caspase 9 after transfection of miR-181 mimic, and the protein expression of cyt C and P53 after transfection of miR-181 inhibitor (Fig. 10 and 11). Therefore, it was speculated that PCP may inhibit TGEV-induced apoptosis through Caspase-dependent mitochondrial pathway after transfection of miR-181.

## 5. Conclusion

PCP possessed the better anti-TGEV effect in vivo. PCP caused the change of miRNA expression profile in the process of inhibiting the infection of PK15 cells by TGEV. PCP could inhibit the replication of TGEV after transfection and apoptosis caused by TGEV by regulating miR-181.

## Author contributions

Xueqin Duan: methodology, investigation, formal analysis, data curation, and visualization, and writing original draft. Nishang Liu and Xingchen Wang: validation. Huicong Li, Xuewen Tan, Yingqiu Liu, Weimin Zhang and Wuren Ma: writing, review, and editing. Yi Wu and Lin Ma: supervision. Yunpeng Fan: Design, supervision, project administration, and funding acquisition. All authors have read and agreed to the published version of the manuscript.

## Acknowledgements

The project was supported by Project funded by National Natural Science Foundation of China (Grant No. 31772788) and National Natural Science Foundation of Shaanxi Province (Grant No. 2020JM-161). We are grateful to all the staff at the Institute of Traditional Chinese Veterinary Medicine of Northwest A&F University for their assistance with these experiments.

